# Sex Differences in Physiological Hair Cycling and Adhesive Material-Induced Hair Regeneration in Mice

**DOI:** 10.64898/2026.07.23.740406

**Authors:** Ying-Ying Chou, Moyuri Inoue, Tomoyuki Toge, Anna Yoshimura, Sachiko Yamashita Takeuchi, Osamu Takahashi, Kazuya Haraguchi, Keisuke Kawasaki, Osamu Kaminuma, Takuma Matsubara, Manabu Habu, Shoichiro Kokabu

## Abstract

Sex differences in hair growth are clinically evident, but sex-dependent regulation of physiological hair cycling and injury-induced hair regeneration remains incompletely understood. We compared physiological dorsal hair-cycle progression and adhesive material-induced localized hair regeneration in male and female C3H/He mice. Males entered the second and third anagen phases earlier than females, indicating longer telogen phases in females. In contrast, localized hair regrowth after application and removal of a cyanoacrylate adhesive material appeared earlier in females. Ovariectomy induced widespread telogen-to-anagen transition and therefore did not permit isolation of ovarian-hormone effects on the localized response. RNA sequencing of intact dorsal skin identified sex-dependent baseline expression profiles involving inflammation, wound response, and tissue repair. Independent time-course quantitative PCR further demonstrated sex-dependent expression of inflammatory and reparative genes after adhesive material application. Local clodronate liposome administration alone induced delayed perifocal hair growth. When combined with adhesive material application, clodronate treatment markedly delayed wound healing and localized hair regrowth in males, whereas these responses were comparatively preserved in females. These findings show that physiological hair cycling and adhesive material-induced hair regeneration exhibit distinct sex differences and suggest that the localized regenerative response is more macrophage-dependent in males than in females.

## Introduction

Sex differences in hair growth and hair cycling are well recognized clinically. One representative example is androgenetic alopecia, which affects both men and women but differs markedly between the sexes in prevalence, age at onset, pattern of hair loss, and clinical progression [1]. These clinical differences have traditionally been attributed to the actions of sex hormones, including androgens and estrogens. However, tissue sex differences also arise from sex-chromosome complement and cell-intrinsic differences in gene expression, epigenetic regulation, and responsiveness to external stimuli [2]. Therefore, understanding sex differences in hair growth and hair cycling requires consideration not only of sex-hormone actions but also of cell- and tissue-intrinsic differences within the skin. Although sex is increasingly recognized as an important biological variable in life-science research [3], cell- and tissue-level sex differences in hair-cycle regulation remain incompletely understood.

Hair follicles are formed during embryonic development and subsequently undergo repeated cycles of anagen, catagen, and telogen throughout life under the control of hair follicle stem cells, producing a new hair shaft during each cycle. The transition of a resting follicle from telogen to anagen is a critical determinant of hair growth. Nevertheless, even in mice, which are widely used in biomedical research, relatively few studies have directly compared physiological hair cycling in males and females over extended periods, and sex differences in telogen duration and the timing of anagen entry remain poorly defined. Estrogen has been reported to maintain murine hair follicles in telogen and suppress anagen entry [4–6]. Conversely, ovariectomy, a widely used approach for investigating ovarian-hormone function, can itself induce a rapid and widespread telogen-to-anagen transition [5]. Thus, in studies using hair cycling or localized hair growth as primary outcomes, systemic hair-cycle changes caused by ovariectomy may obscure the effects of ovarian hormones on a local stimulus. This limitation further emphasizes the need to first compare physiological hair cycling in untreated male and female animals.

Telogen-to-anagen transition can also be induced experimentally by hair plucking, cutaneous wounding, or epidermal injury. In these models, the inflammatory and reparative microenvironment generated after tissue injury contributes to hair follicle stem-cell activation and hair-cycle transition. Cutaneous wound healing itself displays sex differences in inflammatory-cell recruitment, cytokine production, re-epithelialization, and tissue repair [7,8]. Nevertheless, few studies have directly compared wound-induced telogen-to-anagen transition and hair-regrowth kinetics between male and female mice. We recently established a highly reproducible model in which adhesive materials, including pyroxylin and cyanoacrylate, induce localized hair growth following superficial epidermal injury [9]. Although hair regrowth was observed in multiple mouse strains and in both sexes, detailed sex differences in the onset of hair growth, histological progression, and inflammatory and reparative responses were not examined. In addition, our regional analysis of dorsal skin suggested that baseline gene-expression profiles and post-injury inflammatory and reparative responses contribute to differences in the timing of telogen-to-anagen transition among anatomical sites [10]. These observations prompted us to investigate sex differences in adhesive-induced hair regeneration from the perspectives of both the baseline skin state and the response to injury.

In this study, we first compared long-term physiological dorsal hair-cycle progression in male and female C3H/He mice. We then analyzed sex differences in the kinetics and early tissue responses of adhesive material-induced localized hair regeneration. We also examined whether ovariectomy could be used to assess the contribution of ovarian factors in this model. In addition, RNA sequencing of intact dorsal skin and time-course analyses of inflammation- and wound-repair-related genes after adhesive material application were performed to compare baseline and post-treatment skin responses between the sexes. Finally, clodronate liposomes were locally administered to examine the contribution of macrophages to the hair-regrowth response. The aim of this study was to define sex differences in physiological hair cycling and injury-induced telogen-to-anagen transition and to identify associated differences in local inflammatory and reparative responses.

## Materials and Methods

### Animals

Male and female C3H/HeSlc mice were purchased from Japan SLC, Inc. (Hamamatsu, Japan) and maintained under standard laboratory conditions with free access to food and water. Unless otherwise stated, five biologically independent mice were used in each group. All animal experiments were approved by the Experimental Animal Care and Use Committee of Kyushu Dental University (approval number 26-010) and were conducted in accordance with institutional guidelines and the ARRIVE guidelines.

### Observation of physiological hair-cycle progression

Physiological dorsal hair-cycle progression was evaluated in male and female mice. Dorsal hair was shaved with electric clippers under general anesthesia. To observe the transition to the first anagen phase, mice were shaved at postnatal days 22–25 and examined at days 33–36 and 41–44. To assess the subsequent hair cycle, mice were shaved at postnatal days 54–57 and examined at days 128–131, 139– 142, and 199–202. Changes in skin pigmentation and hair regrowth were documented by gross photography.

### Adhesive material-induced localized hair regrowth

Localized hair regrowth was induced by application of a cyanoacrylate adhesive material, as previously described [9,10]. After the dorsal hair was shaved, Aron Alpha Jelly, a gel-type cyanoacrylate adhesive (Toagosei Company, Limited, Tokyo, Japan), was locally applied to the dorsal skin as the adhesive material. After polymerization, the adhesive material was removed together with the attached superficial epidermal layer. The day of adhesive material application was defined as Day 0. Hair regrowth was monitored macroscopically, and skin samples were collected at the indicated time points.

### Ovariectomy

Bilateral ovariectomy or sham surgery was performed in 11-week-old female mice with minor modifications to a previously described procedure [11]. Under general anesthesia, the ovaries were exposed through bilateral dorsolateral incisions. Both ovaries were removed in the ovariectomy group, whereas they were exposed and returned to the abdominal cavity in the sham-operated group. One week after surgery, adhesive material-induced hair regrowth was examined as described above. Successful ovariectomy was confirmed by uterine atrophy at tissue collection [12].

### Histopathological examination

Histological analysis was performed as previously described [9,10]. Skin samples were fixed in 4% paraformaldehyde in phosphate-buffered saline, dehydrated through ethanol and xylene, embedded in paraffin, and sectioned at 4 μm. Sections were stained with hematoxylin and eosin and imaged using a Keyence BZ-X800 microscope (Keyence Corporation, Osaka, Japan).

### Quantitative real-time PCR

Quantitative real-time PCR was performed as previously described [10]. Total RNA was isolated from dorsal skin using a FastGene RNA Basic Kit (Nippon Genetics, Tokyo, Japan) and reverse-transcribed using a High-Capacity cDNA Reverse Transcription Kit (Applied Biosystems, Waltham, MA, USA). PCR was performed using PowerUp SYBR Green Master Mix and a QuantStudio 3 Real-Time PCR System (Thermo Fisher Scientific, Waltham, MA, USA). Skin samples were collected from male and female mice on Days 0, 1, 2, 4, and 6 after adhesive material application. Expression levels of Tnfa, Igf1, Il10, Mrc1, and Il1b were normalized to Tbp using the ΔCT method. Four biologically independent mice were analyzed at each time point, and each sample was measured in technical triplicate. Primer sequences are listed in Supplementary Table 1.

### RNA sequencing and bioinformatic analysis

RNA sequencing was performed using intact dorsal skin from two male and two female mice. These samples were sequenced in the same experiment and analyzed using the same pipeline as in our previous regional analysis of dorsal skin [10]. RNA sequencing and initial bioinformatic analyses were performed by Novogene Co., Ltd. Poly(A) RNA was purified, fragmented, and used for strand-specific library preparation. Libraries were sequenced on an Illumina platform. After quality control, clean reads were aligned to the mouse reference genome mm39 using HISAT2. Gene expression was quantified using raw read counts and FPKM values. Differential expression between female and male skin was analyzed using DESeq2 with adjusted P ≤ 0.05 and |log2 fold change| ≥ 1. Heatmap values are presented as row-scaled z-scores of log2(FPKM + 1).

### Local administration of clodronate liposomes

Clodronate Liposome SMALL and the corresponding control liposomes were obtained from HYGEIA BIOSCIENCE Co., Ltd. (Japan). Immediately before adhesive material application, liposomes were injected locally beneath the planned treatment site using a 29-gauge insulin syringe with a 12.7-mm needle.

For the clodronate group, 25 μL of clodronate liposomes was mixed with 25 μL of control liposomes and administered at a final volume of 50 μL per site. The control group received 50 μL of control liposomes per site. The adhesive material was applied immediately after injection. To assess the effects of liposomes alone, control or clodronate liposomes were also administered without adhesive material application. Gross and histological observations were performed at the indicated time points.

### Statistical analysis

Statistical analyses were performed using GraphPad Prism (GraphPad Software, San Diego, CA, USA). Data are presented as mean ± SD. Quantitative PCR data were analyzed by two-way ANOVA, with sex and time as factors, followed by Sidak’s multiple-comparisons test for comparisons between male and female mice at each time point. P < 0.05 was considered statistically significant.

## Results

### Physiological telogen duration differs between male and female mice

Comparison of dorsal hair cycling between male and female C3H/He mice revealed earlier skin darkening and hair regrowth in males than in females (Figure 1A). This difference was evident during the transition from the first postnatal telogen to the second anagen phase, with a broader area of the dorsal skin entering anagen in males while the corresponding changes were delayed in females.

**Figure 1.**
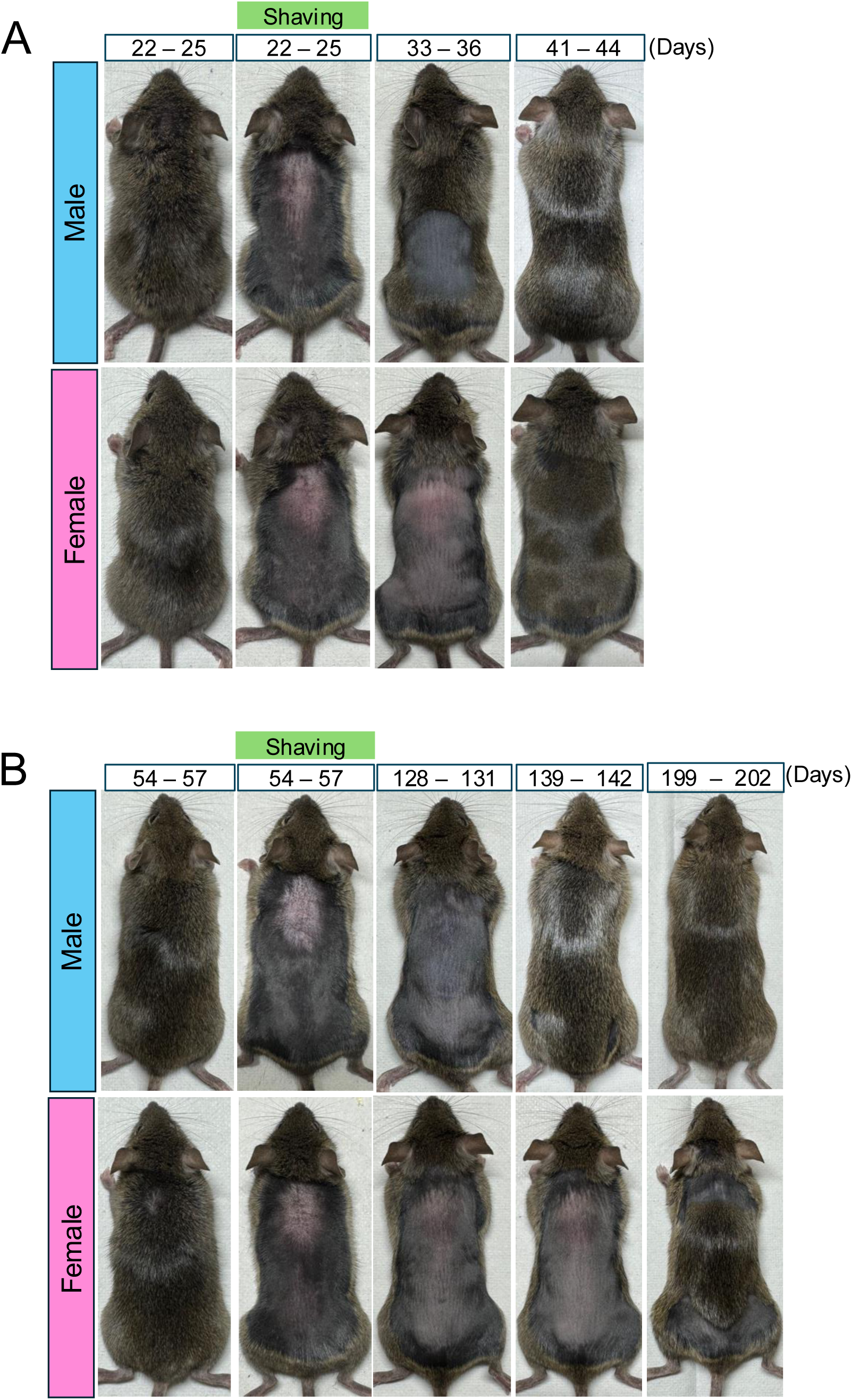
Sex-dependent differences in physiological dorsal hair-cycle progression in C3H/He mice. (A) Sex-dependent differences in dorsal hair-cycle progression during the transition from the first postnatal telogen to the second anagen phase. Dorsal hair of male and female C3H/He mice was shaved at postnatal days 22–25, and subsequent changes in skin pigmentation and hair regrowth were monitored at postnatal days 33–36 and 41–44. Representative dorsal skin images are shown. (B) Sex-dependent differences in dorsal hair-cycle progression from the second anagen phase through the second telogen to the third anagen phase. Dorsal hair of male and female C3H/He mice was shaved at postnatal days 54–57, and subsequent changes in skin pigmentation and hair regrowth were monitored at postnatal days 128– 131, 139–142, and 199–202. Representative dorsal skin images are shown.

A similar sex difference was observed during the subsequent hair cycle. The transition from the second telogen to the third anagen phase, as indicated by changes in skin pigmentation and hair regrowth, occurred earlier in males than in females (Figure 1B). These findings indicate that the timing of telogen-to-anagen transition differs between the sexes and that telogen is longer in female C3H/He mice.

### Adhesive material-induced localized hair regrowth differs between the sexes

Adhesive material application induced localized hair regrowth at the treated sites in both male and female mice; however, the timing and progression of the response differed between the sexes (Figure 2A). Hair regrowth became apparent earlier in females, whereas the response in males was delayed. Histological examination also revealed sex-dependent differences in the progression of wound-associated tissue changes and hair follicle elongation (Figure 2B). Thus, cyanoacrylate induced localized hair regrowth in both sexes, but the kinetics of the response were distinct. To examine whether ovarian factors contributed to this difference, female mice underwent ovariectomy. Successful ovariectomy was confirmed by uterine atrophy (Figure 3A). However, ovariectomized mice exhibited widespread skin darkening and hair regrowth across the dorsal skin rather than a response confined to the adhesive material-treated sites (Figure 3B). Because ovariectomy itself induced a systemic telogen-to-anagen transition, its effect on adhesive material-induced localized hair regrowth could not be evaluated separately in this experimental setting.

**Figure 2.**
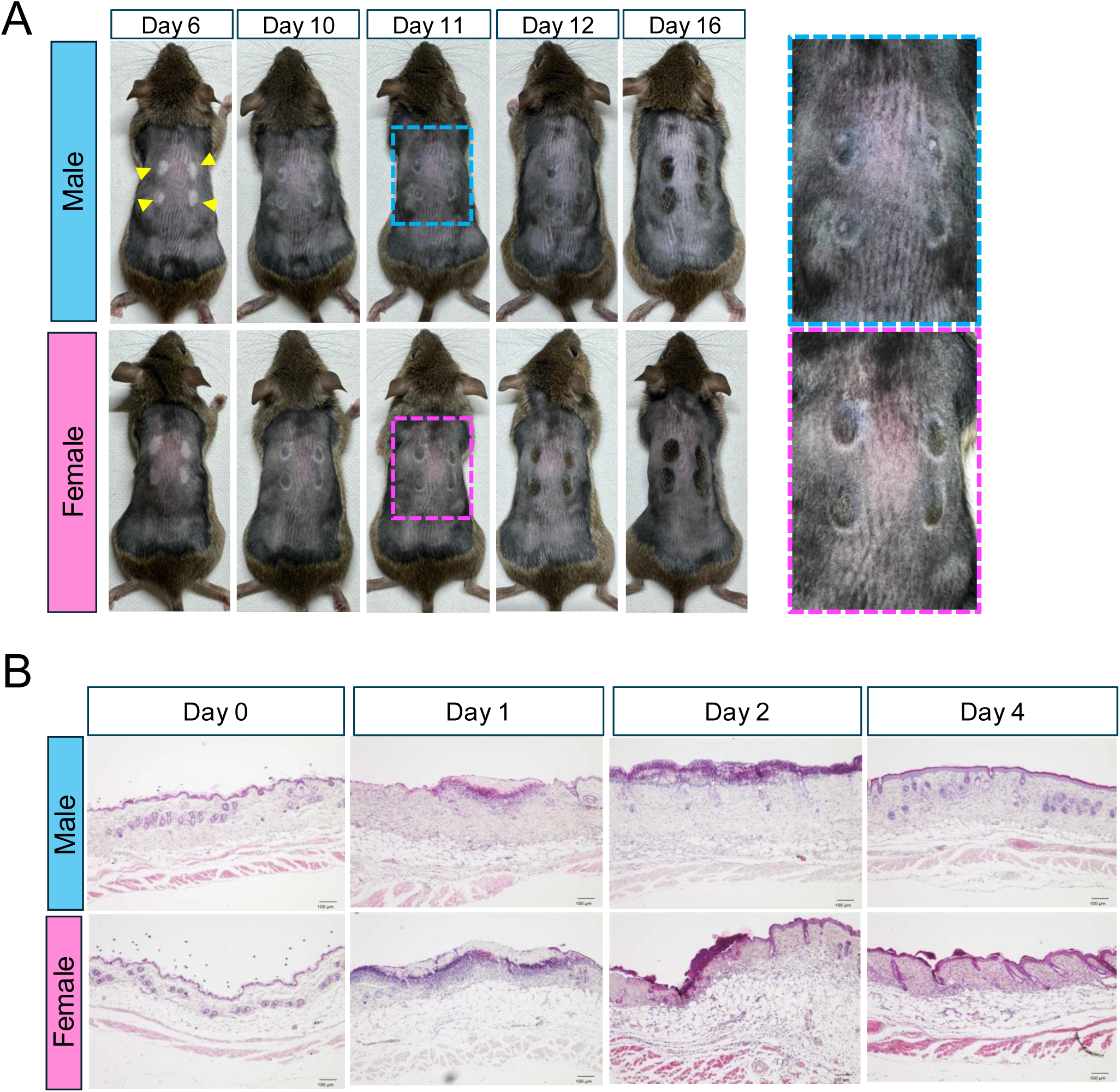
Sex-dependent differences in adhesive material-induced localized hair regeneration. (A) The adhesive material was locally applied to the shaved dorsal skin of male and female C3H/He mice, and subsequent hair-regrowth responses were monitored over time. Representative dorsal skin images on Days 6, 10, 11, 12, and 16 after adhesive material application are shown. Yellow arrowheads indicate adhesive material-treated sites. Representative enlarged images of the boxed regions on Day 11 are shown on the right. (B) Histological analysis of dorsal skin from male and female mice after adhesive material application. Representative hematoxylin and eosin-stained images of skin samples collected on Days 0, 1, 2, and 4 after adhesive material application are shown.

**Figure 3.**
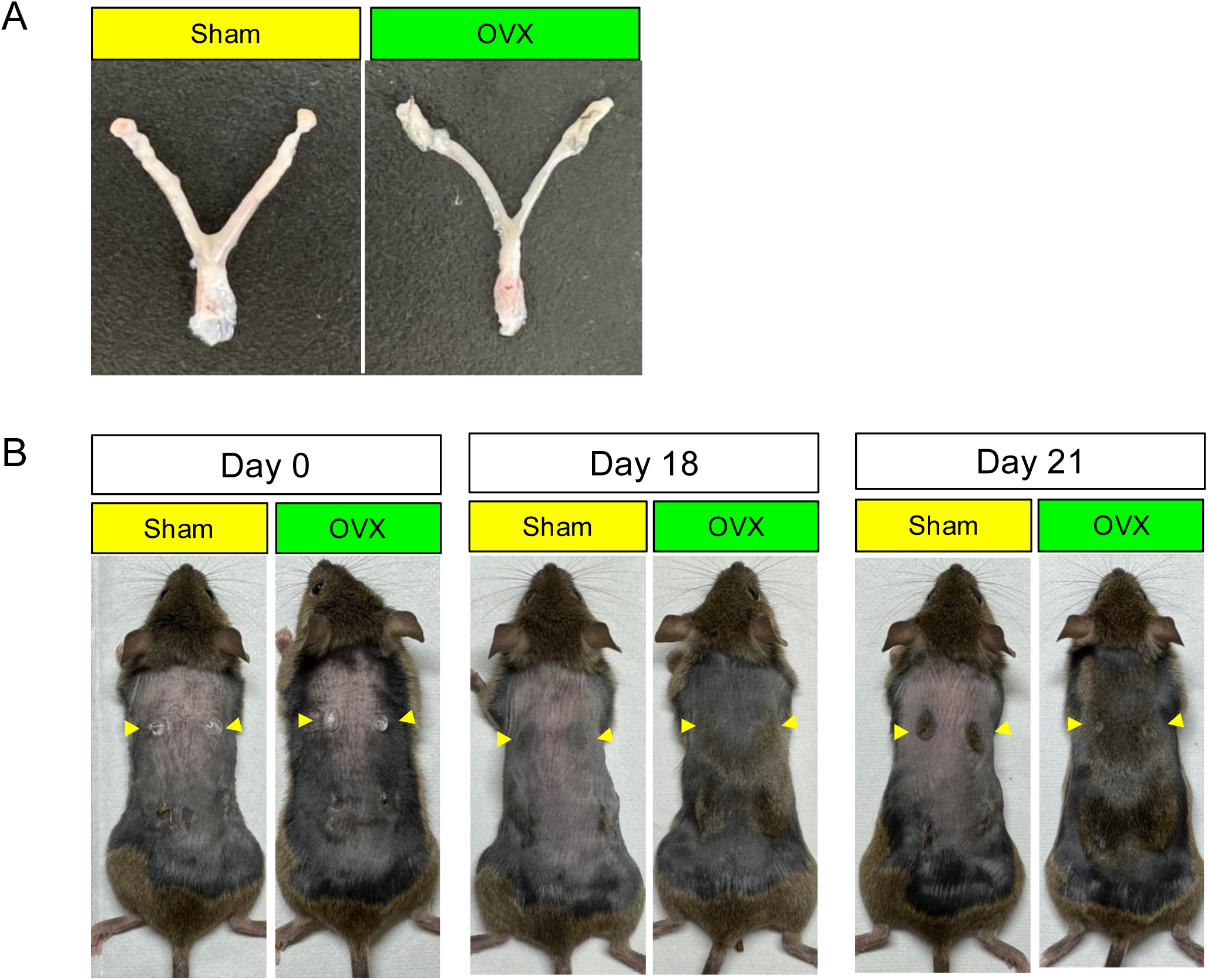
Effects of ovariectomy on adhesive material-induced localized hair regeneration. (A) Representative images of uteri collected after sham operation or ovariectomy. Uterine atrophy in ovariectomized (OVX) mice compared with sham-operated mice confirmed successful ovariectomy. (B) Cyanoacrylate-induced hair-regrowth responses in sham-operated and OVX female C3H/He mice. Sham operation or OVX was performed at 11 weeks of age, and cyanoacrylate was locally applied to the shaved dorsal skin at 12 weeks of age. Representative dorsal skin images on Days 0, 18, and 21 after adhesive material application are shown. Yellow arrowheads indicate adhesive material-treated sites.

### Inflammatory and wound-response programs differ between the sexes in intact and cyanoacrylate-treated skin

To determine whether sex-dependent responses to adhesive material application were associated with pre-existing differences in the skin, gene expression in intact dorsal skin was compared between male and female mice. RNA-seq identified 162 genes expressed at higher levels in females and 123 genes expressed at higher levels in males, demonstrating distinct baseline transcriptional profiles between the sexes (Figure 4A). Differentially expressed genes included Lrrc15, Pla2g2d, Trem2, Cd68, Krt6a, Thbs2, Lyz2, and C3ar1, which are associated with inflammation, wound response, and tissue repair (Figure 4B).

**Figure 4.**
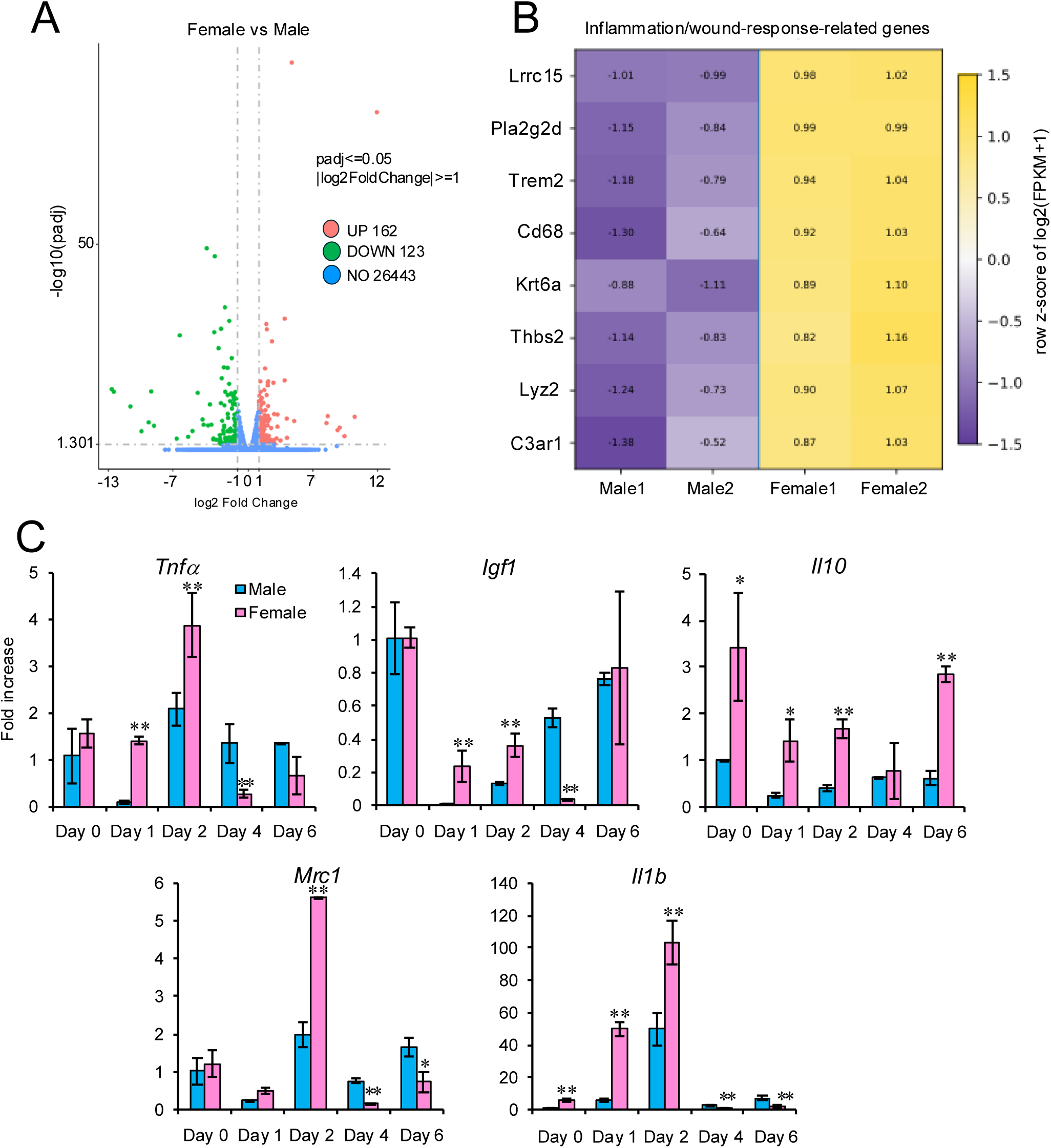
Sex-dependent inflammatory and wound-response gene signatures in intact dorsal skin and after adhesive material application. (A) Volcano plot showing differentially expressed genes between intact dorsal skin samples from female and male C3H/He mice before adhesive material application. Red and green dots indicate genes upregulated and downregulated in female samples relative to male samples, respectively. (B) Heatmap of inflammation- and wound-response-related genes selected from differentially expressed genes identified by RNA-seq analysis of intact dorsal skin. The heatmap is based on the same RNA-seq dataset shown in Figure 4A. Expression values are shown as row-scaled z-scores of log2(FPKM + 1). (C) Quantitative real-time PCR analysis of representative inflammatory and wound-repair-related genes in dorsal skin samples from male and female C3H/He mice after adhesive material application. Skin samples were collected on Days 0, 1, 2, 4, and 6 after adhesive material application, and mRNA levels of Tnfa, Igf1, Il10, Mrc1, and Il1b were examined. Data are presented as mean ± SD. *P < 0.05 and **P < 0.01 for female versus male mice at the same time point.

Time-course analysis after adhesive material application further revealed sex-dependent expression of inflammatory and reparative genes. Il1b and Tnfa were induced during the early response and were expressed at higher levels in females at multiple time points. Il10 and Mrc1 also showed higher expression in females at selected time points, whereas Igf1 displayed a distinct temporal pattern (Figure 4C). These findings indicate that sex differences are present both in the baseline transcriptional state of intact skin and in the inflammatory and reparative response to adhesive material-induced injury.

Local administration of clodronate liposomes alone induced delayed skin darkening and hair regrowth at the injection sites by Day 27 (Figure 5A). When clodronate liposomes were combined with adhesive material application, skin darkening and hair regrowth appeared earlier around the adhesive material-treated sites than after clodronate liposomes alone (Figure 5B), and histological changes consistent with anagen entry were observed (Figure 5C, D). Within the adhesive material-treated area itself, however, clodronate treatment produced a marked sex difference. In male mice, wound closure was substantially delayed, accompanied by delayed hair follicle elongation and localized hair regrowth. In female mice receiving the same dose of clodronate liposomes, wound healing and hair regrowth were comparatively preserved (Figure 5B–D). Because hair follicles surrounding the treated sites responded to clodronate in both sexes, the difference within the wound area was unlikely to result from insufficient local drug action. These findings suggest that adhesive material-induced wound healing and hair regrowth are more highly macrophage-dependent in male than in female mice.

**Figure 5.**
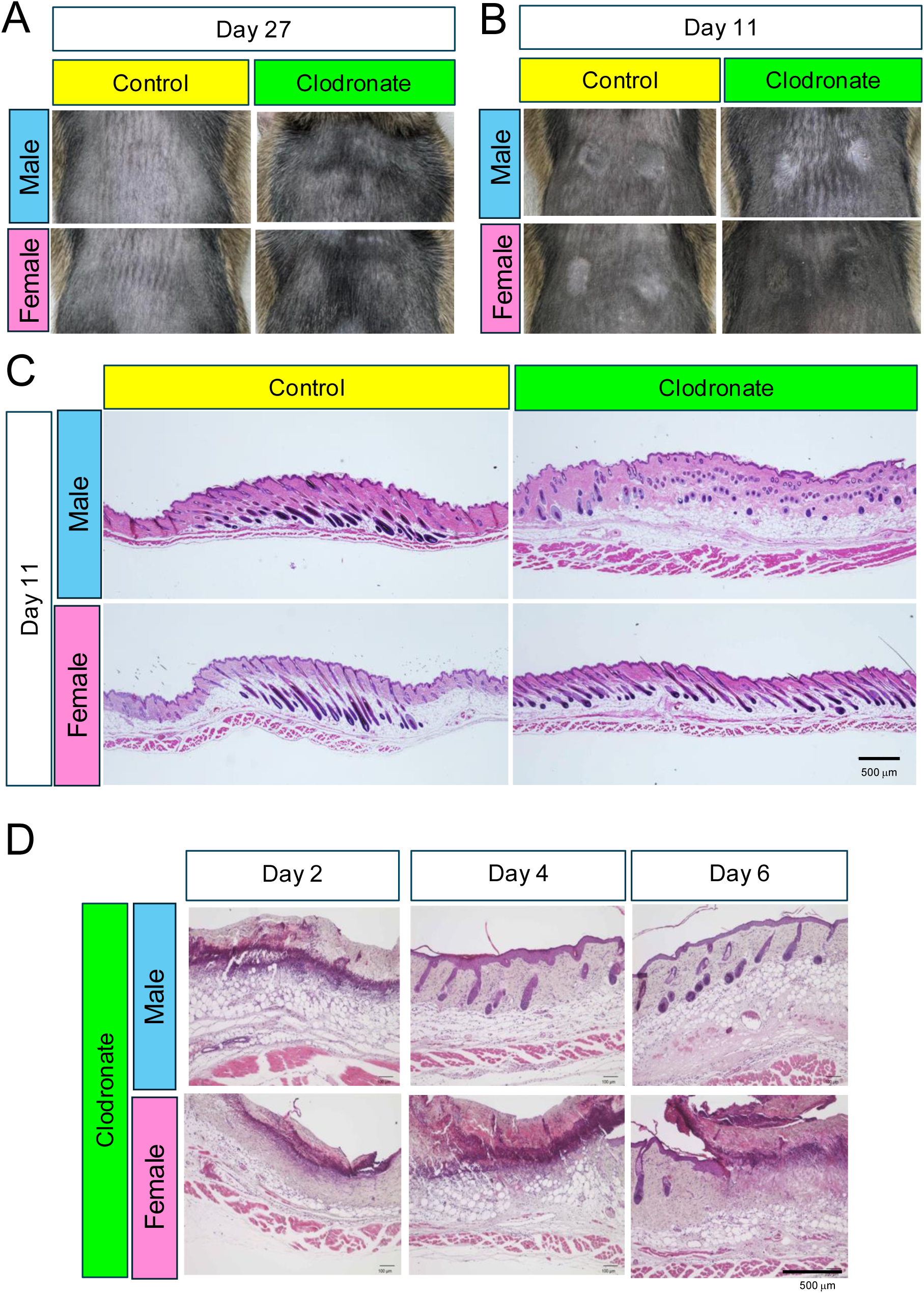
Sex-dependent effects of clodronate liposome treatment on adhesive material-induced localized hair regeneration. (A) Representative dorsal skin images 27 days after local injection of control liposomes alone or clodronate liposomes alone. The adhesive material was not applied. (B) Representative dorsal skin images of male and female C3H/He mice on Day 11 after adhesive material application with local administration of control or clodronate liposomes. (C) Representative hematoxylin and eosin-stained images of dorsal skin from male and female C3H/He mice on Day 11 after adhesive material application with local administration of control or clodronate liposomes. Scale bar, 500 μm. (D) Representative hematoxylin and eosin-stained images of dorsal skin from male and female C3H/He mice on Days 2, 4, and 6 after adhesive material application with local administration of clodronate liposomes. Scale bar, 500 μm.

## Discussion

This study demonstrates sex differences in both physiological hair cycling and adhesive material-induced localized hair-regrowth responses. In C3H/He mice, the transition from the first postnatal telogen to the second anagen phase was delayed in females compared with males, and the second telogen phase also persisted longer in females. After cyanoacrylate application, the timing of hair regrowth, early histological changes, and expression of inflammation- and wound-repair-related genes also differed between the sexes. Baseline gene-expression profiles in intact skin were likewise sexually dimorphic. These findings indicate that sex differences in physiological hair cycling and those in injury-induced localized hair regrowth should be evaluated as distinct biological phenomena.

The sex difference in physiological hair cycling observed here is relevant to the design of mouse hair-cycle studies. In our previous study, 8-week-old female BALB/c, ddY, DBA/2, NC/Nga, C3H/He, and C57BL/6N mice were all in a telogen state permissive for adhesive-induced hair growth [9]. However, it remains unclear which telogen phase is present at that age in each strain, how long it persists, and whether its time course is equivalent in males and females. The present results show that, at least in C3H/He mice, the second telogen phase persisted longer in females than in males.

Ovarian factors, particularly estrogen, are known to influence the hair cycle. Estradiol maintains murine hair follicles in telogen and suppresses anagen entry [4,5,13]. Consistent with previous reports that ovariectomy induces a systemic telogen-to-anagen transition, ovariectomized mice in the present study developed widespread skin darkening and hair regrowth beyond the adhesive material-treated sites. Although ovariectomy is widely used to investigate ovarian-hormone function, it directly alters the primary outcome in studies of hair cycling and localized hair growth. Consequently, the contribution of ovarian factors to adhesive material-induced localized hair regrowth could not be separated from the systemic hair-cycle response to ovariectomy in this model.

Local clodronate liposome administration induced skin darkening and hair regrowth around the adhesive material-treated sites, consistent with the reported activation of hair follicle stem cells by Wnt7b and Wnt10a released during skin macrophage apoptosis [14]. Hair regrowth after clodronate liposomes alone appeared relatively late, whereas the combination of adhesive material application and clodronate treatment produced earlier perifocal darkening and hair growth. One possible explanation is that cyanoacrylate-induced inflammation increased local macrophage recruitment, thereby increasing the number of macrophages undergoing clodronate-induced apoptosis and enhancing hair follicle-activating Wnt signals.

Within the adhesive material-treated area itself, however, clodronate treatment produced a clear sex difference. In males, wound healing was markedly delayed, with corresponding delays in hair follicle elongation and localized hair regrowth. In females treated with the same dose of clodronate liposomes, wound healing and hair regrowth were comparatively preserved. Because hair follicles surrounding the treated sites responded to clodronate in both sexes, this difference is unlikely to be explained by inadequate local drug activity. These findings suggest that adhesive material-induced wound healing and hair regrowth are more highly macrophage-dependent in males than in females. Female skin may retain regenerative capacity through compensatory cell populations or signaling pathways when macrophages are reduced. The RNA-seq analysis in this study used the same sequencing experiment and analytical pipeline as our previous regional analysis of dorsal skin [10], but the shared dataset was analyzed here from the perspective of sex. Major inflammation- and wound-repair-related genes were subsequently examined in independent animals by time-course qPCR. Limitations of this study include the small number of animals used for RNA-seq and the absence of direct measurements of macrophage reduction, apoptosis, and Wnt-pathway activation after clodronate treatment.

In conclusion, sex differences exist in both physiological hair cycling and adhesive material-induced localized hair-regrowth responses. Injury-induced hair-regrowth differences may involve sex-dependent baseline skin states and inflammatory and reparative responses after injury. In addition, adhesive material-induced wound healing and hair regrowth may be more highly macrophage-dependent in males than in females. These findings underscore the importance of considering sex as a biological variable in studies of hair cycling and hair regeneration.

## Author Contributions

Conceptualization: Shoichiro Kokabu.

Data Curation: Ying-Ying Chou, Moyuri Inoue, Tomoyuki Toge, Anna Yoshimura, Sachiko Yamashita Takeuchi.

Formal Analysis: Osamu Takahashi, Kazuya Haraguchi, Keisuke Kawasaki, Ying-Ying Chou, Moyuri Inoue.

Funding Acquisition: Shoichiro Kokabu.

Investigation: Ying-Ying Chou, Moyuri Inoue, Tomoyuki Toge, Anna Yoshimura, Sachiko Yamashita Takeuchi.

Methodology: Ying-Ying Chou, Moyuri Inoue, Tomoyuki Toge, Anna Yoshimura, Sachiko Yamashita Takeuchi, Shoichiro Kokabu.

Project Administration: Shoichiro Kokabu.

Resources: Osamu Kaminuma, Takuma Matsubara, Manabu Habu. Supervision: Shoichiro Kokabu.

Validation: Osamu Takahashi, Kazuya Haraguchi, Keisuke Kawasaki, Osamu Kaminuma, Takuma Matsubara, Manabu Habu.

Visualization: Ying-Ying Chou, Moyuri Inoue, Tomoyuki Toge, Shoichiro Kokabu. Writing – Original Draft: Shoichiro Kokabu.

Writing – Review & Editing: Ying-Ying Chou, Moyuri Inoue, Tomoyuki Toge, Anna Yoshimura, Sachiko Yamashita Takeuchi, Osamu Takahashi, Kazuya Haraguchi, Keisuke Kawasaki, Osamu Kaminuma, Takuma Matsubara, Manabu Habu, Shoichiro Kokabu.

## Data Availability Statement

The RNA sequencing data generated and analyzed in this study will be deposited in the NCBI Gene Expression Omnibus (GEO). The accession number will be provided upon completion of data deposition. All other relevant data are included within the manuscript and its Supporting Information files.

